# Compound Screening with Deep Learning for Neglected Diseases: Leishmaniasis

**DOI:** 10.1101/2021.10.02.462874

**Authors:** Jonathan Smith, Hao Xu, Xinran Li, Laurence Yang, Jahir M. Gutierrez

## Abstract

Deep learning provides a tool for improving screening of candidates for drug re-purposing to treat neglected diseases. We show how a new pipeline can be developed to address the needs of repurposing for Leishmaniasis. In combination with traditional molecular docking techniques, this allows top candidates to be selected and analyzed, including for molecular descriptor similarity.

## 1 Introduction

Each year, approximately 1 million people are diagnosed with the neglected tropical disease known as leishmaniasis. The disease is caused by protozoan parasites of the genus *Leishmania* and it can manifest in one of three forms: visceral, cutaneous and mucocutaneous. Leishmaniasis is estimated to cause the loss of 2.4 million disability adjusted life years and primarily affects people living in countries who do not invest in the development of new treatments (Charlton et al., 2017). Specifically, leishmaniasis is endemic to the Northern Africa, Middle East, Southwest Asia, and Latin America regions.

Current treatments against leishmaniasis not only are difficult to access, but they can also cause adverse side affects and may rely on a non-optimal delivery mechanism (Charlton et al., 2017). Between 2000 and 2017, 0.8% of global investments in infectious disease research went towards studying leishmaniasis, totalling USD 0.8 Billion which ranks Leishmaniasis in 7th place (out of 34) in terms of research spending per disability-adjusted life year among neglected tropical diseases (Head et al., 2020). Despite this, very few drugs have been approved for use against Leishmaniasis, mainly due to the large cost of *de novo* drug development (average of USD $802 million) (Dickson & Gagnon, 2004).

Thus, there is a need to develop new treatments against neglected diseases in a cheaper, safer, and more effective manner. An alternative method for tackling this problem is to repurpose existing drugs used in a different condition or disease. Drug repurposing, also known as repositioning, can significantly reduce the time between compound identification and final deployment by many years while also reducing costs (Li et al., 2016). The goal of drug repurposing is to identify molecules that have either been approved or that underwent clinical trials for another different disease, and identify those molecules with high likelihood of efficacy in treating the target condition. As a matter of fact, a recent success story in this field is the repurposing of Miltefosine, an anticancer agent, as an oral treatment against visceral and cutaneous leishmaniasis (Sunyoto et al., 2018).

In this work, we present a pipeline for drug repurposing targeting leishmaniasis (github.com/jajsmith/drug-repurposing-leishmaniasis). We developed our pipeline as part of our participation at the Indaba Grand Challenge: Curing Leishmanisis hosted by Zindi (zindi.africa). In this challenge, 3D structural files (.pdb, .smf) of both protein targets and candidate drug molecules were provided by the competition organizers. We used these structural files as well as publicly available protein-compound interaction datasets to train multiple machine learning models that rank protein-compound pairs according to their likelihood of interacting. We used the trained models to predict new protein-compound pairs that would be effective at inhibiting target *Leishmania* proteins. Pairs with high predicted binding affinity were then passed through a molecular docking simulation to further assess binding potential and rank them by their predicted affinity. The top three candidates identified by each team were evaluated separately by the competition organizers and the best result was considered as the final scored submission.

Among the top drugs identified by our pipeline, we found Lacosamide (go.drugbank.com/drugs/DB06218) as a potential drug repurposing candidate. Lacosamide is an approved drug for treating partial onset seizures. This drug received a simulated PyRosetta (Chaudhury et al., 2010) docking score of −31.751 against the *Leishmania* enzyme sterol 14-alpha demethylase, an essential component for membrane biogenesis. Thus, there are many promising avenues for experimental validation for this and other protein-drug pairs scored by our pipeline that may contribute to finding novel therapies against leishmaniasis.

## 2 Methods

Drug discovery is a long and involved process with many stages prior to clinical evaluation. Initial research typically focuses on identifying target pathways where the inhibition or activation of a protein will affect the disease before a drug-like molecule or therapeutic can be found through a combination of compound screening, secondary assays and *in vivo* analysis (Hughes et al., 2011). A candidate can then be evaluated for additional constraints like toxicity and preclinical safety, as well as delivery mechanism and cost.

In our work we primarily focus on compound screening. Many protein targets from different *Leishmania* species have been validated already, and we used a set of proteins provided by the challenge organizers as well as public databases of protein-compound interactions were used as training data for binding prediction models. This approach is similar to that described in Dassi et al. (2021) but with a larger set of models analyzed and comparisons between different splits of the provided data based on organism. Below we describe the data used in our work and the models and comparison methods for evaluating binding affinity prediction.

### 2.1 Data

#### 2.1.1 Indaba Grand Challenge Competition Data

The challenge provided three sets of protein targets, a set of all potential targets, a set of preferred targets, and a set of targets found only in *Leishmania major*. There were a total of 103,207 targets provided in FASTA format in the *all targets* set, 8,499 in the *Leishmania major* set and 34,594 in the *preferred* set.

#### 2.1.2 Data for training deep learning models

We experimented with two kinds of models: a multi-objective neural network binding affinity prediction model (MONN) (Li et al., 2020) and the DeepPurpose (Huang et al., 2020) framework. MONN was trained with data from the PDBBind v2018 database (Wang et al., 2005), while DeepPurpose was trained with BindingDB (Gilson et al., 2016).

PDBBind is a collection of 21,382 biomolecular complexes including 17,679 protein-ligand complexes which were used for training MONN.

BindingDB is a database of measured binding affinities. As of July 31, 2021 BindingDB contains 41,300 entries containing 2,303,972 binding data for 8,561 protein targets and 995,797 small molecules.

### 2.2 Evaluation Metrics

#### 2.2.1 Indaba Score

The original competition evaluation of a protein-ligand pair’s binding affinity uses PyRosetta (Chaudhury et al., 2010) to conduct a simulated docking and record the free energy score, which is then used to rank submissions. It is important to note that the exact scoring function used by the challenge organizers to score submissions was not made available to participants upon request until after the competition was over.

#### 2.2.2 Binding metrics (*K*_*d*_, *K*_*i*_, *IC*_50_)

*K*_*d*_, the dissociation constant, is an equilibrium constant that describes the tendency of a complex to break down into smaller pieces. For a general reversible reaction:

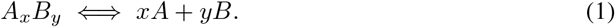

The dissociation constant is defined as:

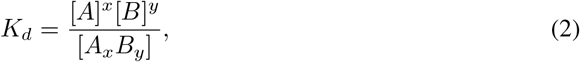

Where [*A*], [*B*] and [*A*_*x*_*B*_*y*_] are the equilibrium concentrations of *A, B*, and *A*_*x*_*B*_*y*_. Therefore, a lower *K*_*d*_ indicates better binding affinity. *K*_*i*_, the inhibitor constant, is a measure of how effective an inhibitor is. *IC*_50_, the half-maximal inhibitory concentration, is an indication of the amount of a substance in inhibiting a specific biological function by half. These two constants can be interconverted for competitive agonists and antagonists using the Cheng-Prusoff equation (Cheng & William, 1973):

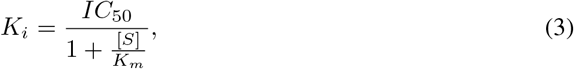

where *K*_*i*_ is the inhibitor binding affinity, *IC*_50_ is the functional strength of the inhibitor, [*S*] is fixed substrate concentration, and *K*_*m*_ is the Michaelis-Menten constant. These three parameters, *K*_*d*_, *K*_*i*_, and *IC*_50_ were chosen as they are provided as affinity measurements in the BindingDB database (Gilson et al., 2016).

### 2.3 Molecular docking and virtual screening

We used AutoDock Vina (Trott & Olson, 2010) version 1.1.2 to dock protein-drug pairs. To automate screening across thousands of candidate pairs, we wrote a pipeline in Python that loads, preprocesses, and prepares protein/drug structural files in PDB format for docking. Briefly, we first use Open Babel (version 2.4) to preprocess PDB files to adjust charges, add hydrogens, and remove water molecules. Then, we use BioPython (Cock et al., 2009) to identify the search box in the protein 3D structure to run docking over. By default, we set the search box volume to the minimum cube that includes the entirety of the protein atoms. Finally, we use AutoDock Vina to dock each protein-drug pair and convert the top predicted pose to a single structural file in PDB format.

### 2.4 Deeppurpose

DeepPurpose (Huang et al., 2020) is a deep learning toolkit for molecular modelling and prediction. We used DeepPurpose to predict the interactions between drug candidates and *Leishmania* protein targets. DeepPurpose uses BindingDB as one of its training datasets in order to train five models with various drug-protein encoder pairs. Each of these models can be applied to a highly customizable classifier for predicting the binding affinity as one of three binding metrics. Among the five pre-trained DeepPurpose models, there are four drug encoders included in the framework: Convolutional Neural Network (CNN) (Krizhevsky et al., 2012), Multi-Layer Perceptrons (MLP) on Morgan (Rogers & Hahn, 2010), Daylight Fingerprint ^1^ and Message Passing Neural Network (MPNN) (Gilmer et al., 2017); as well as two protein encoders: CNN and MLP on Amino Acid Composition (AAC) (Gromiha, 2010) used to train through the BindingDB dataset.

1. **CNN** encodes SMILES and amino acids with an embedding layer. The CNN convolutions are then applied, which is followed by a global max-pooling layer.
2. **Daylight** transforms the SMILES to path-based fingerprints, which is a 2,048-length bits vector containing followed by a multi-layer perceptron.
3. **Morgan** fingerprint is similar to Daylight fingerprint but with a 1,024-length and containing circular radius-2 substructures.
4. **MPNN** takes chemical descriptors as inputs and generates a molecular graph-level embedding vector.
5. **AAC** is an 8,420-length vector characterizing amino acid k-mers, the k-length (k *≤* 3) amino acid subsequences. It also contains the percentage of individual amino acids towards the entire sequence string.

### 2.5 MONN

MONN is a deep learning method for predicting binding affinity between proteins and ligands (Li et al., 2020). It makes further use known 3D binding structures to predict the pairwise non-covalent interactions for a given pair using a CNN encoder for the protein and an MPNN encoder for the molecule.. This provides additional ways to explain predictions using the predicted interactions and the attention layers in the model. The different method also allows us another point of comparison with the final molecular docking simulations. We trained two different models, one for predicting binding affinity on new compounds only, and one for predicting binding affinity on new compounds and new proteins, following the methods described by the original authors. The evaluation performance of these models can be seen in Table 4.

**Table 1:** Pearson Correlations between methods on the top 597 pairs.

| Pearson Correlation | Vina | DeepPurpose | Indaba Score | MONN |
| --- | --- | --- | --- | --- |
| Vina | - | 0.3412 | 0.1936 | 0.3999 |
| DeepPurpose | - | - | 0.5512 | - |
| Indaba Score | - | - | - | 0.3684 |
| MONN | - | - | - | - |

**Table 2:**
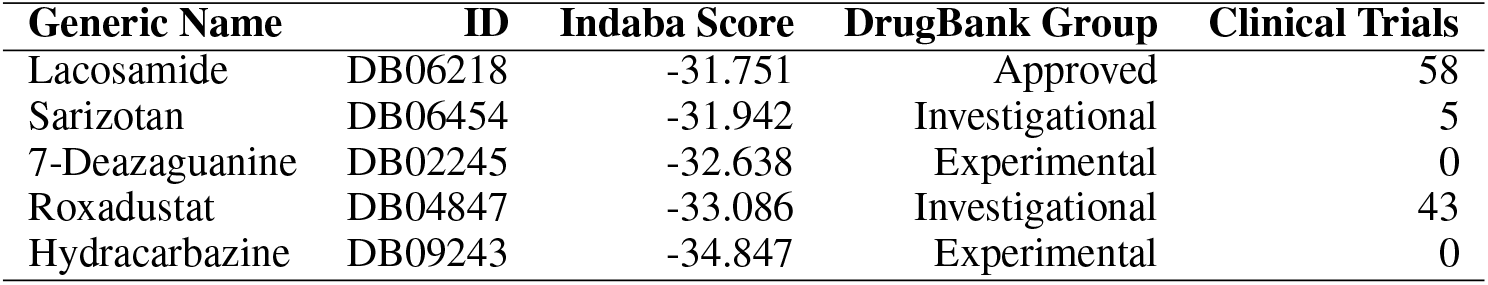
Top 5 drug candidates for the sterol 14-alpha demethylase protein from the DrugBank database along with the number of Completed or Recruiting clinical trials.

**Table 3:** Spearman correlation of predicted binding affinity with label.

| Molecule Encoder<br>Protein Encoder | CNN<br>CNN | Daylight<br>AAC | Morgan<br>AAC | Morgan<br>CNN | MPNN<br>CNN | Sample size |
| --- | --- | --- | --- | --- | --- | --- |
| Leish (Kd) | 0.6713 | 0.7342 | 0.5244 | 0.4056 | -0.0349 | 12 |
| Non-Leish (Kd) | 0.6502 | 0.7077 | 0.6244 | 0.6459 | 0.0950 | 1200 |
| Leish (IC50) | 0.3739 | 0.4593 | 0.0948 | 0.1481 | 0.0895 | 156 |
| Human (IC50) | 0.1617 | 0.4218 | 0.2351 | -0.1707 | 0.3179 | 438 |
| Leish (Ki) | 0.3004 | 0.1688 | 0.2697 | 0.3686 | 0.0455 | 99 |
| Human (Ki) | -0.0613 | -0.0007 | 0.0409 | 0.2007 | -0.0234 | 404 |
| Rand. Non-Leish (Ki) | 0.0765 | 0.1584 | 0.1581 | 0.1819 | 0.0488 | 100 |

**Table 4:** Performance of MONN models with different cross-validation techniques.

| Model | RMSE | Pearson | Spearman | Avg Pairwise AUC |
| --- | --- | --- | --- | --- |
| New Compounds | 1.489 | 0.6893 | 0.6918 | 0.9339 |
| New Compounds + New Proteins | 1.7512 | 0.5220 | 0.5313 | 0.8149 |

Although this approach might seem to leverage more data, since it is making use of the full 3D structure of the protein, it does limit the amount of data that can be used compared with DeepPurpose. The limited data on protein structures for organisms like Leishmania could explain why this approach ends up with lower correlation on the top predicted compound-ligand pairs as seen in Table 1 where the Pearson Correlation of MONN with the Indaba Score 0.1828 less than that of DeepPurpose.

### 2.6 Clustering of molecules by applying PCA of their molecular descriptors

We use the ‘mordred’ (Moriwaki et al., 2018) and RDKit (Landrum, 2019) to generate real-value vector representation for each molecule by 1,613 descriptors. After removing the descriptors whose returns contain any errors and missing values and standardized the vectors, we obtained the molecule feature vector associated with 1,080 descriptors.

The distance matrix among drug candidates is computed based on these features. For each protein, the drugs which are close to the top drugs (i.e. the drugs that have the highest binding affinity with this protein) are selected for further investigation. Then, we use principal component analysis (PCA) to reduce the feature dimension to 3 and visualize molecules to investigate clusters. We further compute the correlation matrix between the Indaba scores and these descriptors.

## 3 Results

### 3.1 Top predicted target-drug pairs

The highest score we received during the Zindi-Indaba competition was ranked 2nd place in the final round of the challenge evaluation. This pair comprised the PubChem substance with identifier SID 56341311, and the *Leishmania infantum*’s protein Sterol 14-alpha demethylase (PDB 3L4D). According to PubChem and ChEMBL, SID 56341311 is a legacy, undefined compound deposited by Thomson Pharma with molecular weight of about 681 Da. This compound does not have any registered clinical trials as of August 31, 2021. This compound, also registered as PF-02575799, was developed by modifying dirlotapide, an FDA-approved drug to treat canine obesity by inhibiting microsomal triglyceride transfer protein. PF-02575799 was developed with improved gut selectivity and had advanced to human phase 1 trials (Robinson et al., 2011).

Among our other top scored candidates, Lacosamide stands out because it is a drug that has already received approval as an oral treatment for partial onset seizures in adults. It has a strong predicted binding score with the sterol 14-alpha demethylase enzyme based on the Indaba Score. Lacosamide is also identified as PubChem substance SID 175427063 and it has favourable predicted ADMET features such as being a non-inhibitor of important human enzyme proteins and low acute toxicity in rats (2.1629 LD50, mol/kg). See it’s structure in Figure 1.

**Figure 1:**
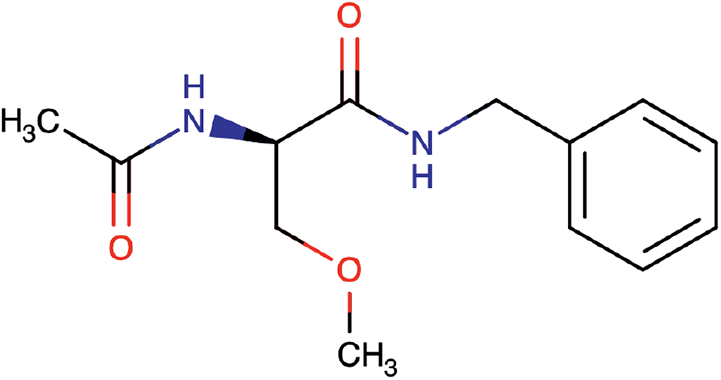
Chemical structure of Lacosamide

Four of the top five DrugBank chemicals are still under investigation or are experimental, and only one other one has had a large number of clinical trials. Drugs with relatively little existing literature will not profer the same speed ups or cost reduction in the development process that drug repurposing is hoping to achieve. Roxadustat (SID 347827699) has completed phase I single-dose and is underingoing phase II US studies as a treatment for Anemia. Predicted ADMET characteristics are not currently available.

As for the target, the sterol 14-alpha demethylase is an enzyme that catalyzes the removal of the 14-alpha-methyl group from sterol precursors. Given that this reaction is essential for membrane biogenesis, this protein has been studied as a potential target for antileishmanial chemotherapy. We also present the top 5 drug candidates for this target in Table 2.

### 3.2 Pre-trained DeepPurpose model selection

As the proteins in the training data are human proteins, we hypothesize that understanding the performance of DeepPurpose across different organisms is crucial to understanding the model’s general applicability. Using protein-chemical pairs from BindingDB which have actual *K*_*d*_*/K*_*i*_*/IC*_50_ values, the pre-trained model performance was evaluated by these values. This was done by (1) electing some BindingDB data in a specific category, (2) running DeepPurpose pre-trained models to generate an affinity rank, and (3) calculating the Spearman correlation between the produced rank and the actual rank. Based on the results as shown in Table 3, Daylight-AAC achieved the highest correlation over different species. For *Leishmania* with a *K*_*d*_ value, the Spearman correlation reached 0.7342, which was the highest among all trials.

### 3.3 Model Comparison

We compared the model predictions against the Indaba Scores for the top 597 submitted protein-compound pairs. The results are shown in Table 1. The MONN model is found to have the highest correlations with Autodock Vina binding scores, while the DeepPurpose Daylight-AAC model has the highest correlations with the Indaba Scores (PyRosetta docking).

### 3.4 Clustering of Molecular Descriptors

We hypothesised that the drugs which best inhibit a given protein target may share similar properties. Molecule clustering based on molecular descriptors is used to select promising subgroups among massive drug candidates. We first investigated the submitted drugs for the Leishmania major pteridine reductase 1 (PDB 6RXC) and found clear clusters for top drugs (7/34) in the three-dimensional plot as seen in Figure A.1 ^2^. Then we computed the distance matrix between the drug candidates in in-trial and drug central database and the top drugs which have an Indaba score smaller than *−*25. The drugs ‘ZINC000012503187’ (Conivaptan) and ‘ZINC000100013130’ (Midostaurin) are very close, and they are both top drugs for *Leishmania donovani* lanosterol 14-alpha-demethylase (UniProtKB E9BAU8) as seen in Figure A.2 ^3^.

We further explored the molecular properties which may affect the binding affinity between molecules and a given protein. According to the correlation matrix between Indaba score and 1080 molecular descriptors ^4^, the ‘VSA Estate6’ and ‘Estate VSA7’ descriptors show the highest correlation with the Indaba score with the correlation of −0.31 and −0.29 respectively. They are all Hybrid EState-VSA descriptors. We also present the top 10 descriptors in Table A.1.

## 4 Conclusions

In summary, we identified several drug candidates for treatment of Leishmaniasis by applying deep learning to the task of compound screening for drug repurposing. We compared multiple models and final scoring evaluations as part of the Indaba Grand Challenge: Curing Leishmaniasis. Finally we compare our top-scoring candidates through a clustering analysis of their molecular properties. Further research can be done in the form of *in vivo* analysis to determine the efficacy of this candidate and any other toxicity, safety, and delivery constraints.

This work has indicated that there is still room for exploration in drug repurposing techniques to address neglected diseases. Future work can expand the approach presented here to other neglected diseases. It is our hope that deep learning models can contribute to the global fight against infectious diseases.

## Author Contributions

All authors worked on the competition submissions and wrote this manuscript. HX, XL, LY developed the DeepPurpose model selection and candidates as well as running the molecular descriptor clustering analysis. JS tuned the MONN model and conducted analysis of molecular docking and model scoring comparisons. JG wrote the docking pipeline.

## Acknowledgments

This work was supported by Queen’s University and the Natural Sciences and Engineering Research Council of Canada (NSERC) [RGPIN-2020-06325].

## A APPENDIX A: Clustering analysis of top drug-protein pairs

**Figure A.1:**
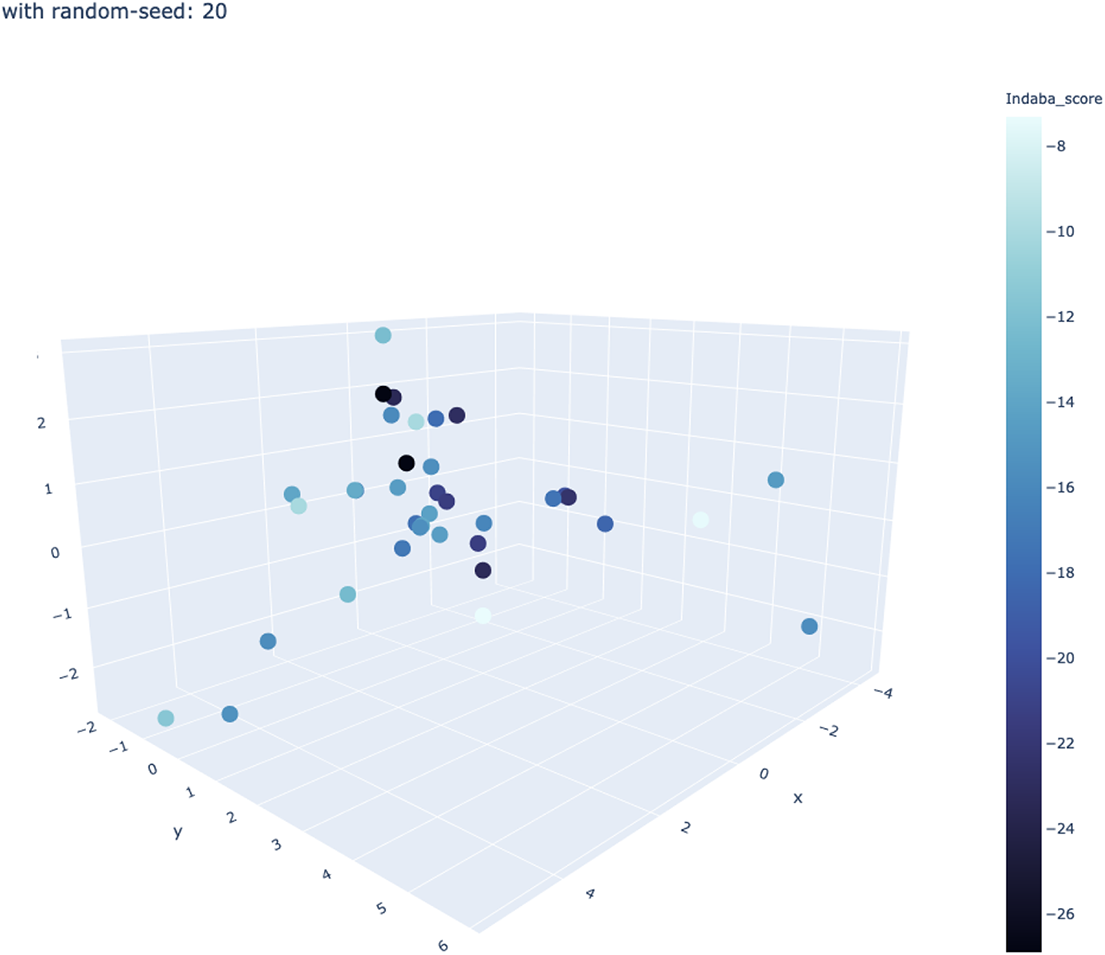
Clusters of top drugs for protein 6RXC by Indaba score

**Figure A.2:**
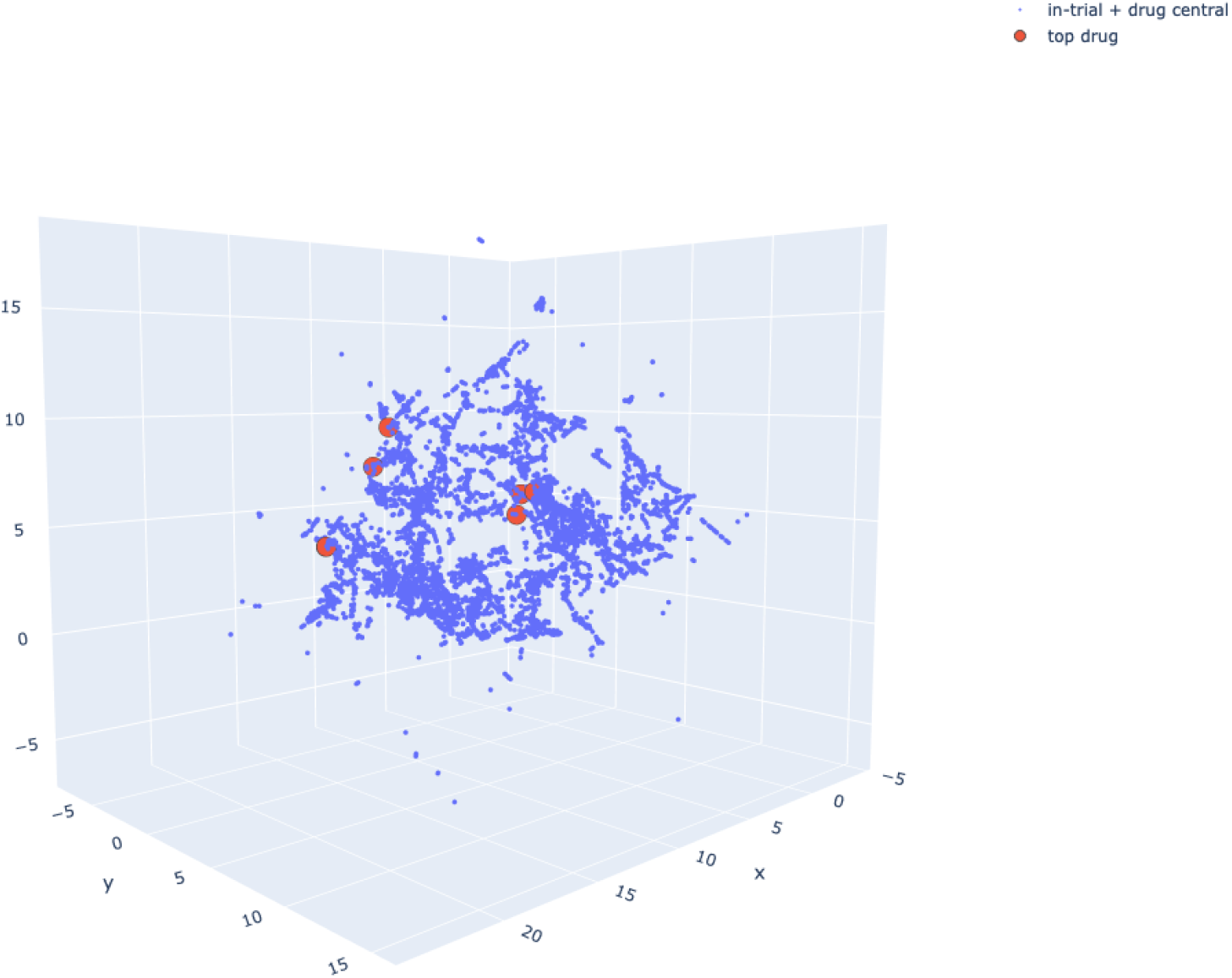
Clusters of top drugs from the in-trial and drug central databases

**Table A.1:**
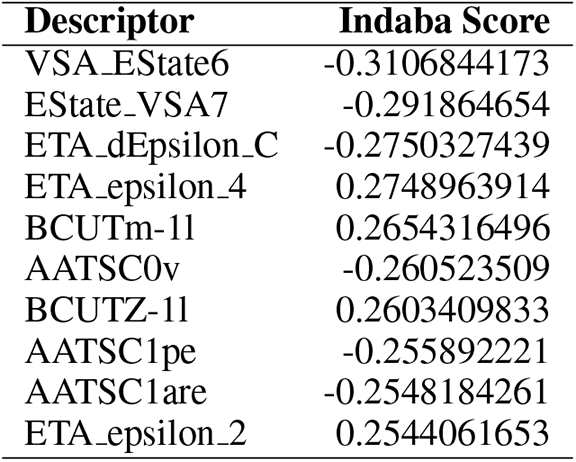
Top 10 descriptors correlated to Indaba Score.

## Footnotes

1 Daylight Theory Fingerprints: https://www.daylight.com/dayhtml/doc/theory/theory.finger.html

2 https://drive.google.com/file/d/1em6dU6bjfl8Qk3ekpKCGw6bFpeYn2Th_/view?usp=sharing

3 https://drive.google.com/file/d/1GD2FqAIjibyTlyW9tDLkGs2cObidqe4a/view?usp=sharing

4 The full correlation matrix is available at: https://docs.google.com/spreadsheets/d/1CFWxCtPfBmQsOGFVtYHTuHiAXYe7xrKAOlrd03_ybSQ/edit?usp=sharing

## References

Rebecca L. Charlton, Bartira Rossi-Bergmann, Paul W. Denny, and Patrick G. Steel. Repurposing as a strategy for the discovery of new anti-leishmanials: the-state-of-the-art. Parasitology, 145, 2: 219–236, August 2017. doi: 10.1017/S0031182017000993. URL https://www.ncbi.nlm.nih.gov/pmc/articles/PMC5964475/.

Sidhartha Chaudhury, Sergey Lyskov, and Jeffrey J. Gray. Pyrosetta: a script-based interface for implementing molecular modeling algorithms using rosetta. Bioinformatics, 26 5:689–91, 2010.

Yung-Chi Cheng and Prusoff H. William. Relationship between the inhibition constant (ki) and the concentration of inhibitor which causes 50 per cent inhibition (i50) of an enzymatic reaction. Biochemical Pharmacology, 22(23):3099–3108, 1973.

Peter J. A. Cock, Tiago Antao, Jeffrey T. Chang, Brad A. Chapman, Cymon J. Cox, Andrew Dalke, Iddo Friedberg, Thomas Hamelryck, Frank Kauff, Bartek Wilczynski, and Michiel J. L. de Hoon. Biopython: freely available Python tools for computational molecular biology and bioinformatics. Bioinformatics, 25(11):1422–1423, 03 2009. ISSN 1367-4803. doi: 10.1093/bioinformatics/btp163. URL https://doi.org/10.1093/bioinformatics/btp163.

Loic Dassi, Hassan Kane, and Ebenezer Nkwate. Computationally accelerating protein-ligand docking for neglected tropical diseases: A case study on drug repurposing for leishmaniasis. ICLR 2021 Workshop: Machine Learning for Preventing and Combating Pandemics, 2021.

M. Dickson and J. P. Gagnon. The cost of new drug discovery and development. Discov Med, 4 (22):172–179, Jun 2004.

Justin Gilmer, Samuel S Schoenholz, Patrick F Riley, Oriol Vinyals, and George E Dahl. Neural message passing for quantum chemistry. In International conference on machine learning, pp. 1263–1272. PMLR, 2017.

Michael K Gilson, Tiqing Liu, Michael Baitaluk, George Nicola, Linda Hwang, and Jenny Chong. Bindingdb in 2015: a public database for medicinal chemistry, computational chemistry and systems pharmacology. Nucleic acids research, 44(D1):D1045–D1053, 2016.

M. Michael Gromiha. Chapter 2 - protein sequence analysis. In M. Michael Gromiha (ed.), Protein Bioinformatics, pp. 29–62. Academic Press, 2010. ISBN 978-81-312-2297-3. doi: 10.1016/B978-8-1312-2297-3.50002-3. URL https://www.sciencedirect.com/science/article/pii/B9788131222973500023.

Michael G Head, Rebecca J Brown, Marie-Louise Newell, J Anthony G Scott, James Batchelor, and Rifat Atun. The allocation of us 105 billion in global funding for infectious disease research between 2000 and 2017: An analysis of investments from funders in the g20 countries. Available at SSRN 3552831, 2020.

Kexin Huang, Tianfan Fu, Lucas M Glass, Marinka Zitnik, Cao Xiao, and Jimeng Sun. Deeppurpose: a deep learning library for drug–target interaction prediction. Bioinformatics, 36(22-23): 5545–5547, 2020.

J. P. Hughes, S. Rees, S. B. Kalindjian, and K. L. Philpott. Principles of early drug discovery. British Journal of Pharmacology, 162(6):1239–1249, 03 2011. doi: 10.1111/j.1476-5381.2010.01127.x. URL https://www.ncbi.nlm.nih.gov/pmc/articles/PMC3058157/.

Alex Krizhevsky, Ilya Sutskever, and Geoffrey E Hinton. ImageNet classification with deep convolutional neural networks. In Advances in Neural Information Processing Systems, volume 25. Curran Associates, Inc., 2012. URL https://proceedings.neurips.cc/paper/2012/hash/c399862d3b9d6b76c8436e924a68c45b-Abstract.html.

G Landrum. Rdkit: Open-source cheminformatics. https://wrdkit.org/. Accessed, 10, 2019.

Jiao Li, Si Zheng, Bin Chen, Atul J. Butte, S. Joshua Swamidass, and Zhiyong Lu. A survey of current trends in computational drug repositioning. Briefings in bioinformatics, 17(1):2–12, 2016. doi: 10.1093/bib/bbv020. URL https://www.ncbi.nlm.nih.gov/pmc/articles/PMC4719067/.

Shuya Li, Fangping Wan, Hantao Shu, Tao Jiang, Dan Zhao, and Jianyang Zeng. Monn: a multiobjective neural network for predicting compound-protein interactions and affinities. Cell Systems, 10(4):308–322, 2020.

Hirotomo Moriwaki, Yu-Shi Tian, Norihito Kawashita, and Tatsuya Takagi. Mordred: a molecular descriptor calculator. Journal of cheminformatics, 10(1):1–14, 2018.

Ralph P Robinson, Jeremy A Bartlett, Peter Bertinato, Andrew J Bessire, Judith Cosgrove, Patrick M Foley, Tara B Manion, Martha L Minich, Brenda Ramos, Matthew R Reese, et al. Discovery of microsomal triglyceride transfer protein (mtp) inhibitors with potential for decreased active metabolite load compared to dirlotapide. Bioorganic & medicinal chemistry letters, 21(14):4150–4154, 2011.

David Rogers and Mathew Hahn. Extended-connectivity fingerprints. 50(5):742–754, 2010. ISSN 1549-9596, 1549-960X. doi: 10.1021/ci100050t. URL https://pubs.acs.org/doi/10.1021/ci100050t.

Temmy Sunyoto, Julien Potet, and Marleen Boelaert. Why miltefosine—a life-saving drug for leishmaniasis—is unavailable to people who need it the most. BMJ Global Health, 3(3), 2018. doi: 10.1136/bmjgh-2018-000709. URL https://gh.bmj.com/content/3/3/e000709.

Oleg Trott and Arthur J. Olson. Trott o, olson aj. autodock vina: improving the speed and accuracy of docking with a new scoring function, efficient optimization, and multithreading. J Comput Chem., 31(2):455–461, 2010. doi: 10.1002/jcc.21334. URL https://www.ncbi.nlm.nih.gov/pmc/articles/PMC4719067/.

Renxiao Wang, Xueliang Fang, Yipin Lu, Chao-Yie Yang, and Shaomeng Wang. The pdbbind database: methodologies and updates. Journal of medicinal chemistry, 48(12):4111–4119, 2005.

